# EnTAP: Bringing Faster and Smarter Functional Annotation to Non-Model Eukaryotic Transcriptomes

**DOI:** 10.1101/307868

**Authors:** Alexander J. Hart, Samuel Ginzburg, Muyang (Sam) Xu, Cera R. Fisher, Nasim Rahmatpour, Jeffry B. Mitton, Robin Paul, Jill L. Wegrzyn

**Affiliations:** Department of Ecology and Evolutionary Biology, University of Connecticut, Storrs, CT, USA; Department of Ecology and Evolutionary Biology, University of Colorado Boulder, Boulder, CO, USA 80309

**Keywords:** Transcriptomics, Gene Expression, Functional annotation, Non-model, Genome annotation

## Abstract

EnTAP (Eukaryotic Non-Model Transcriptome Annotation Pipeline) was designed to improve the accuracy, speed, and flexibility of functional gene annotation for *de novo* assembled transcriptomes in non-model eukaryotes. This software package addresses the fragmentation and related assembly issues that result in inflated transcript estimates and poor annotation rates, while focusing primarily on protein-coding transcripts. Following filters applied through assessment of true expression and frame selection, open-source tools are leveraged to functionally annotate the translated proteins. Downstream features include fast similarity search across three repositories, protein domain assignment, orthologous gene family assessment, and Gene Ontology term assignment. The final annotation integrates across multiple databases and selects an optimal assignment from a combination of weighted metrics describing similarity search score, taxonomic relationship, and informativeness. Researchers have the option to include additional filters to identify and remove contaminants, identify associated pathways, and prepare the transcripts for enrichment analysis. This fully featured pipeline is easy to install, configure, and runs significantly faster than comparable annotation packages. EnTAP is optimized to generate extensive functional information for the gene space of organisms with limited or poorly characterized genomic resources.

## INTRODUCTION

While the genomics era has enabled tremendous progress in characterizing new genomes, we have sampled a comparatively small portion of the organismal biodiversity. Among eukaryotes, nearly 4,000 species have been sequenced but fewer than 400 are assembled to the chromosome level, and less than 40 are considered complete [1]. This disparity speaks volumes to the challenges associated with high-throughput, short-read sequences which remain the dominant input for many genome sequencing endeavors. With limited whole genome resources, in terms of species representation and completion, the focus on transcriptomics remains widespread. Transcriptomics, as a subcategory within functional genomics, focuses on quantifying expression levels of the coding region. This measurement can be evaluated in response to abiotic or biotic stimuli, including differences between tissues, developmental stages, or conditions. Techniques focused on assessing the gene space can often provide more insight for specific biological questions for a fraction of the time and cost associated with generating a full reference genome [2].

High-throughput RNA sequencing, commonly known as RNA-Seq, utilizes deep sequencing of short-reads to quantify expression differences [3]. The most widely adopted protocol relies on fragmentation of mRNA into short fragments which are converted to cDNA and processed to prepare a sequencing library [4]. The highly sensitive nature of RNA requires a more robust experimental design with numerous technical and biological replicates [5]. A variety of bioinformatic pipelines have been developed with an emphasis on normalization and quantification to attempt to separate relevant signals from the background noise [6]. Since well resolved reference genomes are seldom available, the *de novo* transcriptome assembly strategy allows us to leverage the redundancy of short-read sequencing to find overlaps between the reads and assemble them into transcripts. One of the most popular transcriptome assemblers, Trinity, traverses the De Bruijn graph to assemble each isoform [7]. The inherent challenges of assembling these short-reads leads to chimeric sequences, fragmented transcripts, and erroneous contigs [8]. It is not unusual to generate a very large number of transcripts, often 3 to 4 times greater, than the estimated gene space for the assessed organism. In addition, the average or N50 values presented for the assembled transcripts are often less than 1Kbp which is attributed to fragmentation. These problems persist despite thoughtful experimental design and minimal to no sample pooling. Despite these challenges, the *de novo* assembled transcriptome represents an important milestone for a previously uncharacterized species and a basis for examining new biological phenomenon [5].

Following RNA-Seq analysis, the assembled transcripts are divided into lists of differentially expressed (DE) genes and remain associated with an arbitrary identifier generated by the *de novo* assembler before they undergo functional annotation. Assignment of the differentially expressed genes to a functional assessment is often more tedious and complex than the process of sequencing and assembly. The ability to efficiently and accurately characterize these transcripts is impacted by the quality of the assembly as well as the magnitude of existing resources in public databases. Several functional annotation pipelines have been developed with the goal of easing the burden for non-expert researchers. The most widely used pipelines, Blast2GO [9] and Trinotate [10], include a combination of sequence similarity with downstream methods that integrate protein domain, gene family, Gene Ontology (GO) terms, and biological pathway assignments. These pipelines are broadly applicable but suffer from runtimes exceeding days or even weeks depending on the databases selected and the HPC resources available. In addition, they operate in a naive fashion without context for species relevance, informativeness of the alignments, or potential for library contamination.

Here, we present a novel, open-source annotation pipeline designed to remedy specific shortcomings of existing packages. EnTAP incorporates efficient database search methods (DIAMOND) with a multi-database approach to improve speed and accuracy [11]. EnTAP implements an alternative approach for selecting the best homology-based alignment. While coverage alignment scores are an important criteria for this selection, the scores are ignorant of phylogenetic relevance, informativeness, and the possibility that other organisms (contaminants) are present in the assembled transcriptome. In EnTAP, these criteria are considered and this information is integrated with relevant and rapid gene family annotations, which provides more context for non-model systems with limited database resources. The combination of speed, accuracy, open-source code, and simple parameterization provides a reliable and flexible platform for the functional annotation of non-model transcriptomes.

## OVERVIEW

EnTAP was developed to contend with the downstream annotation challenges of short-read or long-read, *de novo* transcriptome assembly and provide meaningful functional assignment up to 50 times faster than the current alternatives. EnTAP eliminates the need for web applications, licensed software, as well as many of the challenges of the homology-based transfer approach that can propagate incorrect annotations [12]. This Unix application is designed for simple installation, configuration, and execution. It combines downstream transcriptome filtering and annotation steps while remaining independent of any specific *de novo* transcriptome assembler. It provides realistic parameters and flexibility to customize the analysis to the organism of interest. The execution takes place with a single command that will initiate both phases: *transcriptome filtering* and *transcriptome annotation* (Figure 1).

**Figure 1.**
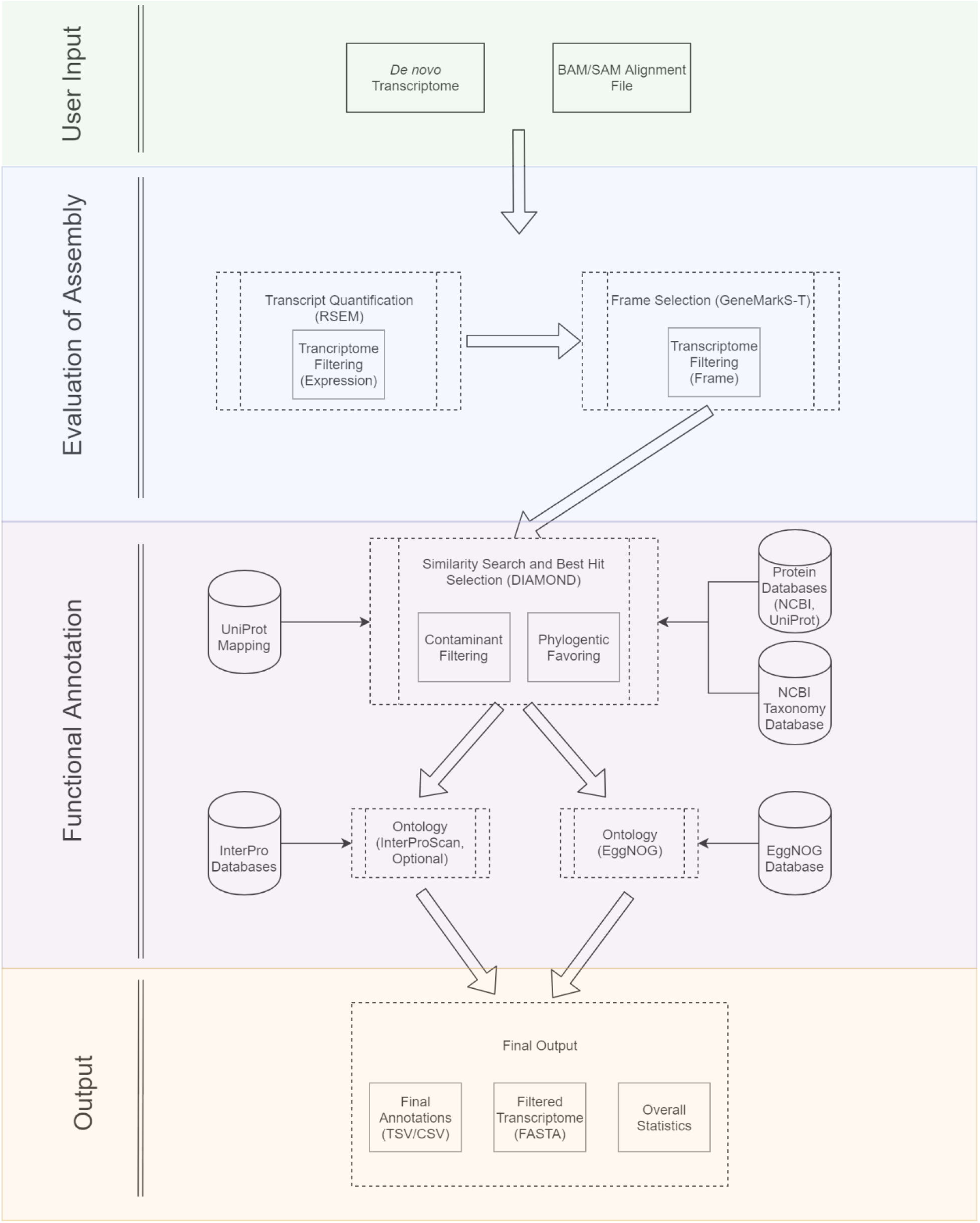
Overview of EnTAP Annotation Pipeline

*Transcriptome filtering* contends with known challenges of fragmentation as well as problematic assemblies. This is represented as two distinct steps in EnTAP, frame selection and expression quantification. Assuming the user provides an optional short-read alignment file, EnTAP will leverage RSEM [13] to remove transcripts that do not meet minimal mapping threshold represented as a normalized FPKM (Fragments Per Kilobase Million). The package is designed to quantify transcript abundance from *de novo* transcriptomes and is applied here to improve the quality of the transcriptome. If this stage is assessed outside of EnTAP, or the user chooses to skip this phase, the transcriptome will move directly to frame selection. In the case of long-read assembled transcriptomes, the assembly would also be directly subject to frame selection as the first step. Frame selection is implemented with GeneMarkS-T which will predict the most likely Open Reading Frame (ORF) or coding region. It will also remove transcripts where no frame was detected and provide a set of trimmed (free of Untranslated Regions (UTR)) nucleotide sequences and translated protein sequences. EnTAP proceeds to the primary annotation steps with the translated and filtered protein sequences.

*Transcriptome annotation* comprises homology (similarity searching), gene family assignment, and GO term/pathway assignment. This stage is designed for protein-coding transcripts. NCBI’s BLAST (Basic Local Alignment Search Tool) is the colloquial application used for the homology stage of functional annotation; although, software such as DIAMOND [11] and RAPSearch2 [14] outperform BLAST in terms of speed, while maintaining similar accuracy. DIAMOND executed in the *sensitive* alignment mode reports speeds up to 2500 times faster than that of BLASTX while reporting 94% of the same matches detected by BLASTX [11]. EnTAP leverages DIAMOND for aligning protein queries (following frame selection) against up to *five* different user-defined protein sequence databases. EnTAP can uniquely detect and process the headers for databases sourced from EMBL-EBI as well as NCBI which provides exceptional flexibility to the user in the selection of the target databases. It is recommended that users select a combination of curated databases (at least 3) that represent full-length proteins. For organisms with very few genomic resources in these databases, it is recommended to include the comprehensive but less curated, nr database (NCBI). The default setting in EnTAP generates a total of three alignments (if available) for each query protein. Oftentimes, these sequences are reported with varying degrees of E-value (measure of the alignment significance), query coverage (percent of query aligning to target sequence from the database), and informativeness. EnTAP offers a unique method of selecting the optimal alignment that factors in the E-value, coverage, taxonomic relevance, contaminant status, and curation of the alignment.

During the configuration step, contaminant taxons, such as *bacteria* or *fungi*, as well as the taxon of the transcriptome itself can be indicated. It is common for RNA-Seq studies to introduce small to moderate levels of RNA that can assemble into full or partial genes from associated organisms [15]. Alignments closer in taxonomic relevance to the user’s species will be favored and contaminants will be unfavored. This phylogenetic filter is made possible by the provision of an origin species for reference protein sequences from curated databases, such as NCBI RefSeq [16] and Uniprot Swiss-Prot [17]. EnTAP will cross-reference this information with NCBI’s Taxonomy Database, and determine the lineage of the origin species. The level of curation, or “informativeness,” is the final consideration in the selection of the optimal alignment. EnTAP utilizes a list of terms, whether provided by the user or default, which will flag an alignment as *uninformative* or *informative.* Descriptive terms such as “predicted” and “unknown” are un-favored. EnTAP will choose the optimal alignment that is closest in lineage to the target species as well as the most *informative* (Figure S1).

Following selection of the optimal target sequence, independent gene family assignment is initiated with a local EggNOG database via EggNOG-mapper [18]. The current release, version 5.0, consisting of 4.4M orthologous groups derived from 379 taxonomic levels, provides an alternative means of Gene Ontology (GO), pathway, and protein domain assignment [19]. Orthologous genes, or genes resulting from speciation, provide a more reliable means of functional annotation for organisms with limited resources. Although a CAFA benchmark has yet to be performed for the GO prediction algorithm, the continual updates of the database, scalability, and improved performance in OrthoBench2 benchmarks provide for a powerful tool to integrate into EnTAP [19, 20]. Alternative approaches for GO assignment, including PANNZER2 [21], Argot2 [22], and others exist although report slower speeds, do not provide for command line integration and may rely heavily on the curated databases such as UniProt The EggNOG associated software, EggNOG-mapper, leverages DIAMOND and SQLITE functionality and has been reported to assign an average of 32 more GO terms per sequence with speeds up to 2.5 times faster than that of InterProScan [18]. GO terms represent a community curated, controlled vocabulary that simplifies information exchange and summary. GO contends with the variable nomenclature used in sequence descriptors by providing terms for three categories: biological function, molecular process and cellular component [23].

EnTAP provides integrated summaries, statistical information, and graphical representations of the data. Metrics such as N50, N90, and longest and shortest sequence lengths are calculated on the provided transcriptome and after subsequent filtering stages. Additional graphical representations are provided to show the distribution of transcripts removed and those remaining. *Transcriptome annotation* includes graphical representations of: taxonomic distribution of contaminants and optimal hits, Gene Ontology term distribution, and EggNOG taxonomic distribution. The final annotation summary is comprehensive and simple to interrogate, as well as parse. The plain text delimited document provides full details for each transcript, including: frame, sequence similarity results, contaminant status, Gene Ontology terms, pathway information, and gene family/protein domain assignment. Intermediate summaries and a comprehensive log file provide additional statistics and tracking information. The full pipeline integrates efficient tools and processes to functionally annotate NGS derived transcriptomes in a fraction of the time of existing platforms.

## EVALUATING FUNCTIONAL ANNOTATION PIPELINES

Several pipelines exist for the functional annotation of transcriptomes that offer varying degrees of speed, accuracy, usability, and computing platforms (Table 1). Blast2GO [9], Trinotate [10], and Annocript [24] can be seen as among the most popular with Google Scholar results (as of June 2019) reporting 13.3K, 784, and 71 citations, respectively. Trinotate, Annocript, and Blast2GO (with a paid subscription) offer annotation through a Unix-based environment while Blast2GO is also accessible via a standalone application. Most services incorporate the traditional NCBI BLAST approach to similarity searching. Trinotate, and Annocript provide open reading frame detection methods that do not require a reference genome which is ideal for *de novo* assembled transcriptomes. Annocript is focused on identification of long non-coding RNAs and limits sequence similarity to well curated UniProt/UniRef databases. Trinotate and Blast2GO Pro were chosen for comparison as the most widely used and comprehensive functional annotation pipelines that are applicable to all eukaryotic organisms. Similar to EnTAP, they integrate across multiple sources with a similar goal of providing a final comprehensive annotation. Trinotate offers a similar annotation method and environment to EnTAP, while Blast2GO Pro provides annotation through an intuitive standalone application and a Unix-based option (available for paid subscribers).

**Table 1.**
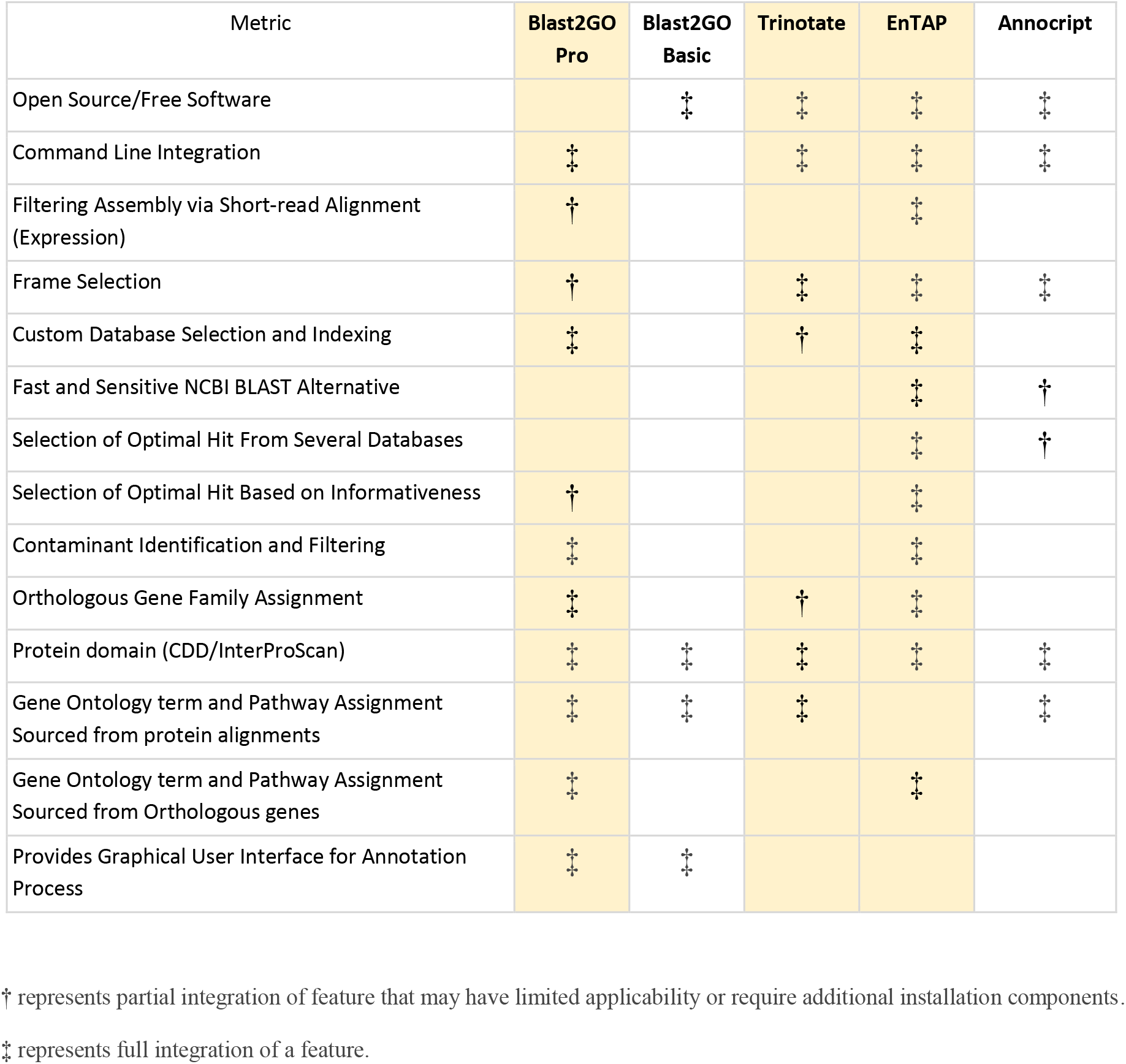
Qualitative Comparison of Functional Annotation Software.

Three non-model, Illumina (paired-end) sequenced and *de novo* assembled transcriptomes were acquired for evaluation with EnTAP, Blast2GO, and Trinotate ranging from 28,350 to 38,640 transcripts. The three species chosen were the *Entylia carinata* (camelback treehopper), *Funaria hygrometrica* (cord-moss), and *Pinus flexilis* (limber pine) providing a varied taxonomic source. The *P.flexilis* and *F.hygrometrica* libraries represent single genotype RNA extractions, while *carinata* is pooled.

### Transcriptome Filtering

Transcriptome filtering was evaluated for frame selection between EnTAP and Trinotate with equivalent stages not examined for Blast2GO since this functionality is not available in the base installation. EnTAP utilizes GeneMarkS-T, while Trinotate executes Transdecoder as a means of frame selection. GeneMARKS-T can process any transcriptome assembly (not assembler specific) while Transdecoder leverages information on Trinity labeled isoforms annotated in the header generated by this assembler. Both methods will remove sequences that do not provide a detectable frame as well as indicate whether the trimmed sequences appear to be full-length, partial (containing 5’ or 3’ ends), or internal (neither end). Both methods will also generate a final peptide sequence.

EnTAP’s implementation of GeneMarkS-T consistently produced more sequences and more complete sequences, while having a slightly lower average sequence length and N50 compared to Trinotate’s Transdecoder (Figure 2A, Table S1). The results of *F. hygrometrica* are similar with frame selection yielding an average sequence length (bp) of 808.79 compared with 761.39 through GeneMarkS-T, and 7213 complete sequences compared to 9695, respectively (Figure 2B, Table S1). Additionally, GeneMarkS-T detected more partial 3’, and fewer internal and partial 5’ sequences compared to Transdecoder. Across the three transcriptomes, EnTAP detected the reading frame on an average of 14% more sequences with a lower overall average sequence length (bp) of 5% and lower N50 by 4% (Table S2, Table S3).

**Figure 2.**
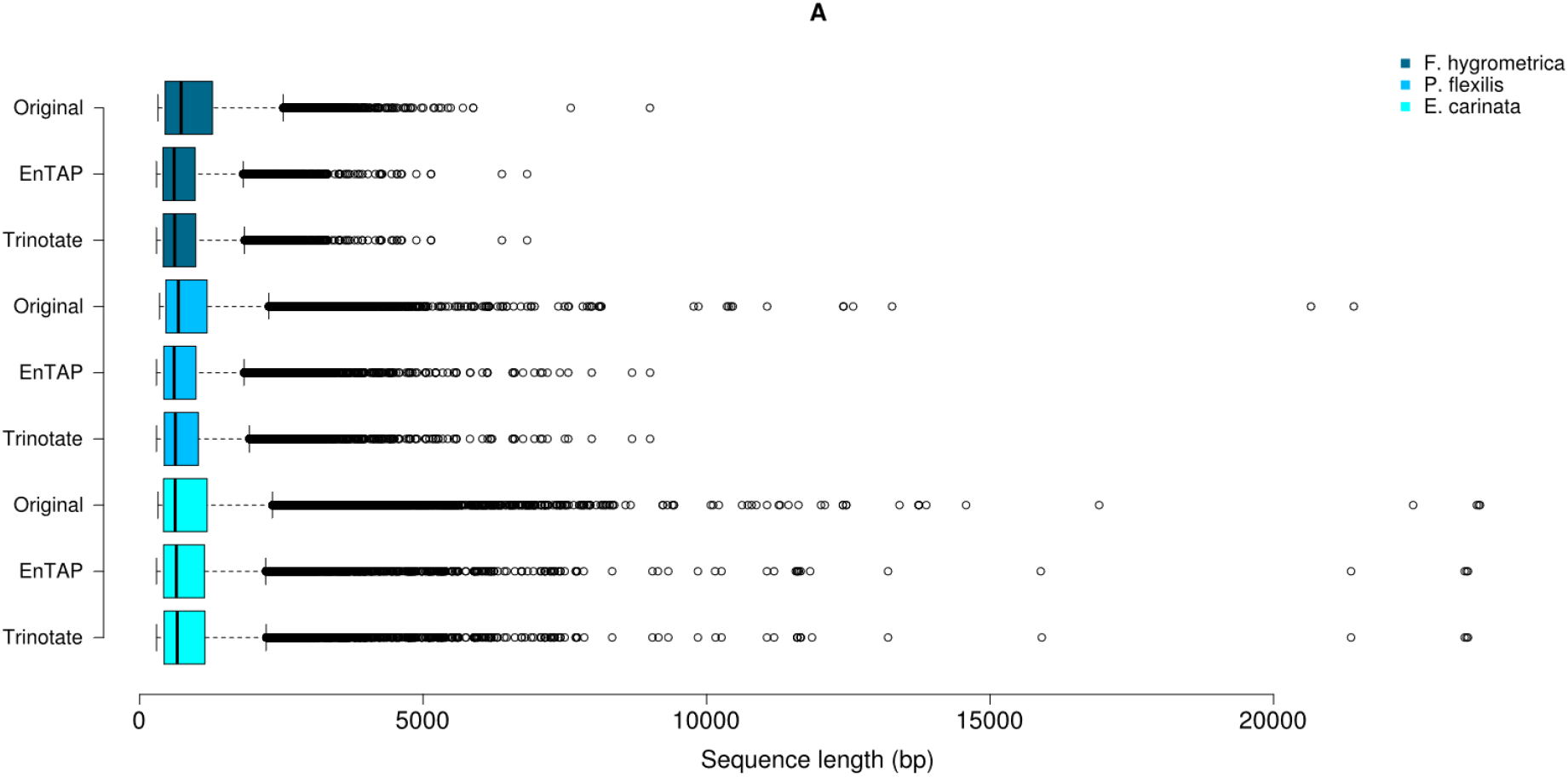
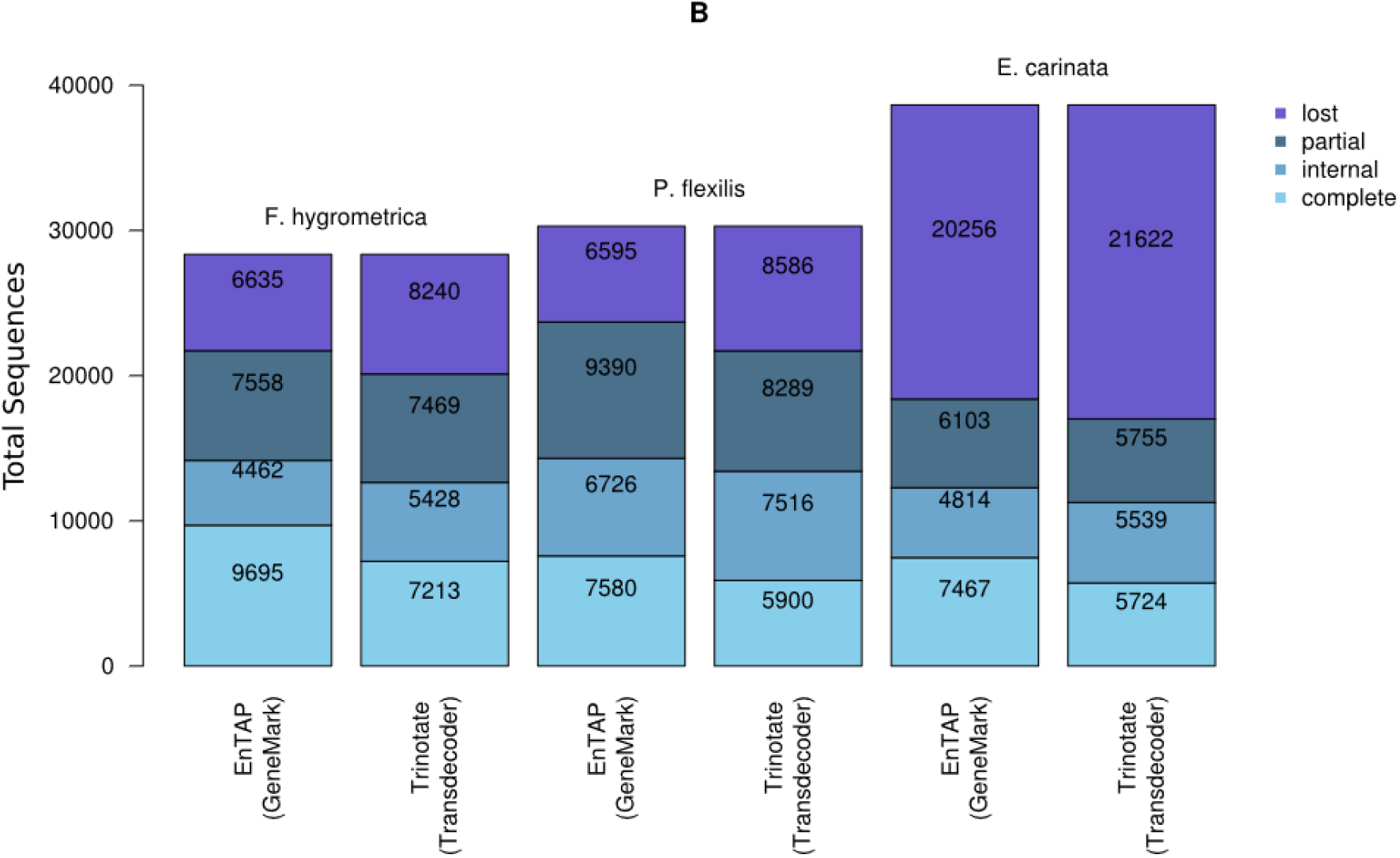
EnTAP and Trinotate Frame Selection Results. (A) Comparison of sequence lengths following the frame selection process between the three species examined across each of the pipeline’s method of frame selection. EnTAP incorporates GeneMarkS-T, while Trinotate utilizes Transdecoder. (B) Comparison of sequence completeness following frame selection.

Further transcriptome filtering can be performed by coupling the optional expression filtering feature (utilizing RSEM) to the transcriptome before the annotation stages of the pipeline. This will remove sequences that do not meet a minimum mapping threshold, providing a higher quality transcriptome downstream. A comprehensive analysis was not performed for this feature across the three pipelines. However, results for the *F. hygrometrica* and *E. carinata* indicated that expression filtering can be helpful in removing sequences that ultimately would be unannotated depending on assembly methods, such as pooled versus non-pooled RNA libraries. This was pronounced in the pooled *E. carinata* results, where 90.85% of the sequences removed (940 sequences) through expression filtering were ultimately not annotated, while the non-pooled *F. hygrometrica* had 65.40% of the sequences ultimately annotating, of the 1448 sequences that were removed.

### Transcriptome Annotation

#### Comparison Across Independent Database Sources

A comparison of homology, or similarity searching, was performed for each of the pipelines based upon completeness, contaminant detection, and phylogenetic relevance of alignments across both pooled and non-pooled transcriptomes. Trinotate utilized NCBI BLAST (BLASTX and BLASTP) functionality, while EnTAP incorporated DIAMOND (BLASTP), and Blast2GO Pro homology searching was performed through NCBI’s CloudBlast (BLASTX), on the paid access package. Similarity searching was initially executed with a minimum query coverage of 50% and minimum E-value of 1E-5 against two curated databases: NCBI’s RefSeq and EMBL-EBI’s Swiss-Prot. In regards to contaminant detection and phylogenetic relevance of EnTAP and Trinotate, optimal alignments were selected through EnTAP’s own methodology and the lowest E-value, respectively.

Trinotate retained the highest alignment rate as a percentage of overall sequences with both EnTAP and Trinotate maintaining similar rates and Blast2GO aligning the fewest (Figure 3A, Figure 3C). This remained true for independent runs against both target databases (RefSeq and Swiss-Prot). *F. hygrometrica* produced alignments against NCBI’s RefSeq for Trinotate, Blast2GO, and EnTAP at 73.24%, 48.99%, and 68.67%, respectively. Alignments against Swiss-Prot resulted in a similar pattern with Trinotate, Blast2GO, and EnTAP seeing 51.94%, 31.73%, and 47.76%, respectively (Table S4, Table S5, Table S6). Blast2GO had lower alignments across each of the three species, hovering around 30% to 50% of the total transcriptome. Given the non-model status of all three organisms, a larger percentage of the transcriptomes are expected to annotate with the more comprehensive RefSeq database (Figure 3C). In addition, it is expected that the majority of the optimal hits assigned from Swiss-Prot are informative since this is a well curated repository. When comparing the results for RefSeq, EnTAP consistently assigned more informative sequences as a result of its selection method (Figure 3A, 3C). The slightly higher percentage of alignment reflected by Trinotate primarily results from inclusion of non-frame selected input sequences for analysis with BLASTX. Of the sequences not detected by EnTAP during this process, upwards of 9% (1908 alignments with *P. flexilis*) were purely BLASTX alignments.

**Figure 3.**
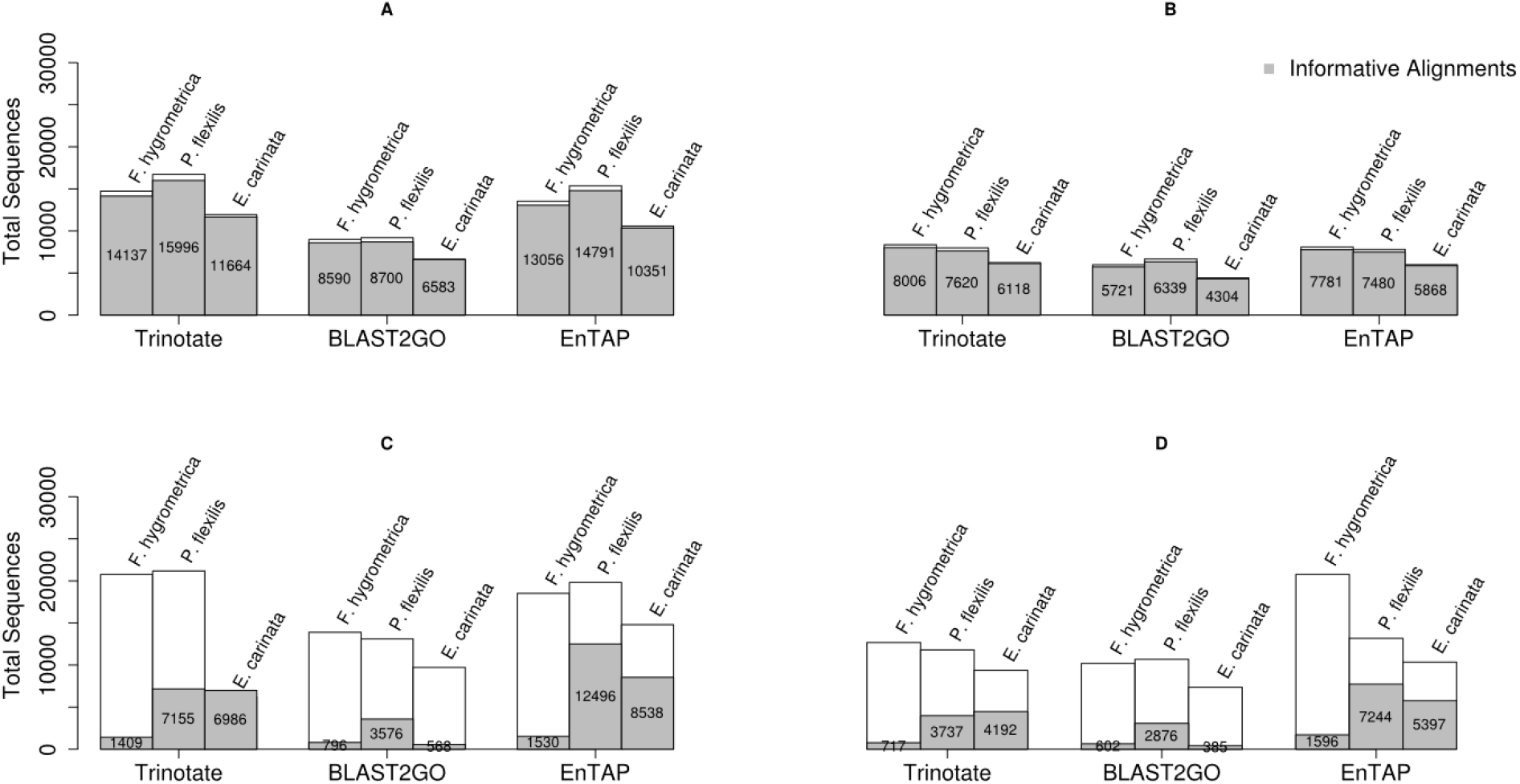
Evaluation of Independent Similarity Search Results – UniProt Swiss-Prot (A,B) and NCBI RefSeq Complete (C,D). (B,C) Homology results when applying both 50% query and 50% target coverage minimum thresholds.

#### Overall Annotation (Independent Sequence Similarity and Gene Family)

Non-model species with limited database resources benefit from functional identification through comprehensive and well annotated orthologous databases that extend beyond protein domain identification. The number of sequences with an annotation is defined here as an alignment resulting from similarity searching or annotation through EggNOG (EnTAP), HMMER against PFam (Trinotate), or InterProScan against PFam (Blast2GO). Furthermore, comparisons were made to encompass unique sequences with a Gene Ontology term assigned and/or KEGG assignments (pathways) from either similarity searches or gene family/protein domain assignment.

EnTAP retained the highest overall annotation rate across all species for Swiss-Prot/EggNOG with Trinotate producing the lowest annotation rate (Swiss-Prot/PFam) (Figure 4A). For example, comparisons with *F. hygrometrica* generated an annotation rate among Trinotate, Blast2GO, and EnTAP, of: 51.94%, 53.11%, and 68.14%, respectively (Table S8). These results can largely be attributed to the inclusion of the EggNOG-mapper approach. Trinotate retained more KEGG pathway and Gene Ontology term assignments for Swiss-Prot with percentages of 45.51% and 50.74% compared to EnTAP’s 22.75% and 43.19%. The assignment of pathways and ontology terms in Trinotate incurs a heavy reliance on Swiss-Prot resulting in lower Gene Ontology term and KEGG pathway assignments for other databases. This can be seen in the RefSeq results, where EnTAP produced the highest overall Gene Ontology and KEGG pathway assignment when compared to Trinotate and Blast2GO. Trinotate (RefSeq/PFam) provided a higher overall annotation compared to EnTAP (RefSeq/HMMER) (Trinotate: 73.23%, EnTAP: 71.74%) and a much higher annotation rate when compared to Blast2GO (RefSeq/InterProScan) (61.81%) (Figure 4B). The reliance of Trinotate on the Swiss-Prot database does not allow for pathway annotation from other databases with the exception of ontology terms from PFam. In the *F. hygrometrica* against RefSeq example, EnTAP assigned a KEGG pathway term to 22.75% of sequences, with Blast2GO and Trinotate annotation generating 9.92% and 0%, respectively.

**Figure 4.**
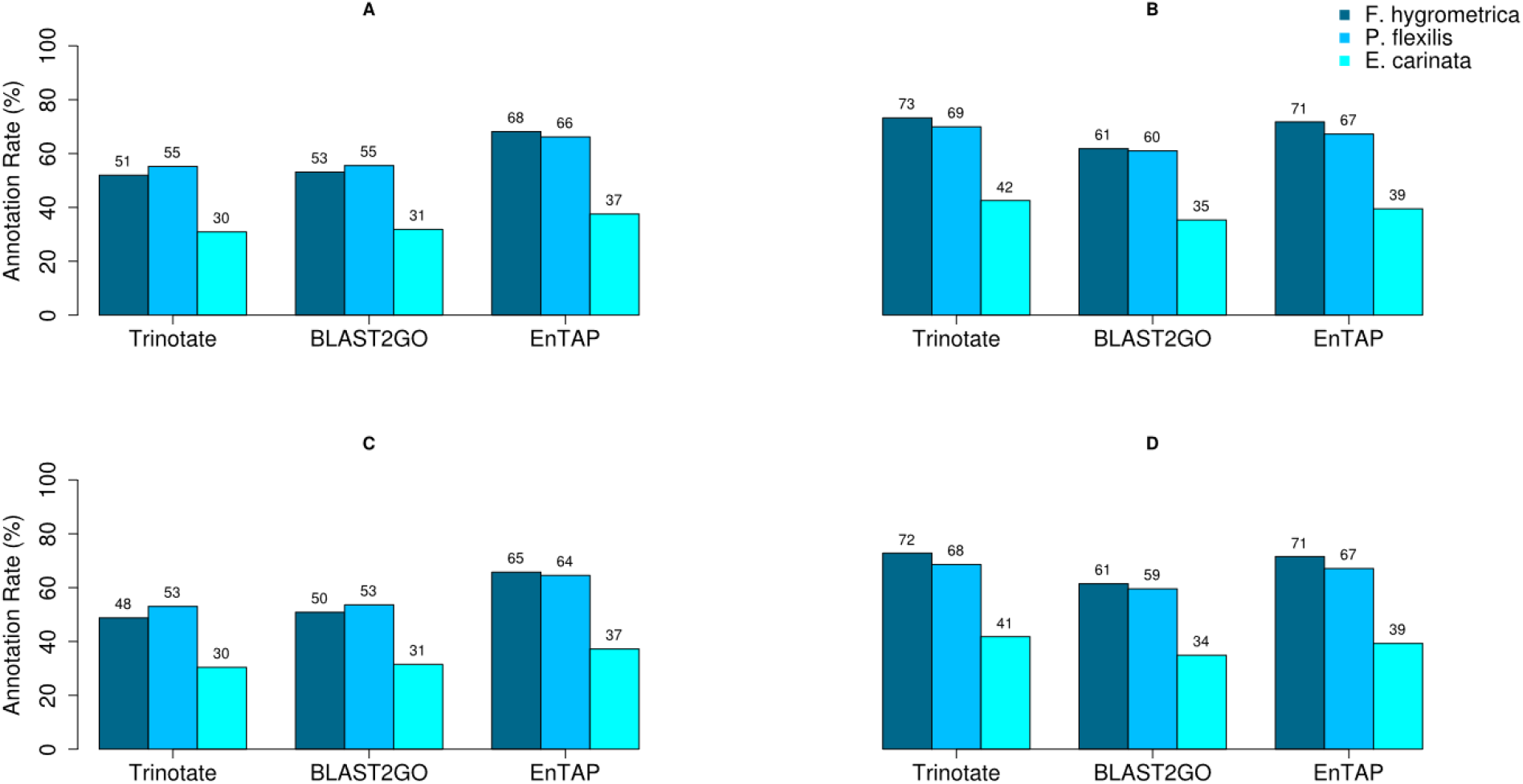
Overall Annotation Rate – UniProt Swiss-Prot (A,C) and NCBI RefSeq Complete (B,D). (C,D) Annotation results with the removal of bacterial and fungal contaminants.

The slight annotation disparity between EnTAP and Trinotate against RefSeq decreased when contaminant removal was applied to the results (Figure 4C, Figure 4D). When considering the results of *F. hygrometrica*, the disparity in annotation between EnTAP and Trinotate decreased from 71.74% and 73.23% to 71.53% and 72.83%, respectively. Across the three species, annotation against the RefSeq database brought EnTAP and Trinotate’s annotation within 1-2%, while EnTAP continued to report a higher overall annotation rate against the Swiss-Prot database. Removing contaminants from the results also had the effect of reducing the Gene Ontology percentage differences between EnTAP and Trinotate from 3-6% for the Swiss-Prot database, and bringing Blast2GO and EnTAP within 1-2% of each other when considering overall annotation against the RefSeq database.

#### Evaluating Quality of the Annotation

EnTAP’s algorithm includes a minimum query and target coverage of 50%. Since other annotation programs only allow modification of the query coverage, the target percentage parameter was not included in the initial comparisons. To examine the impact of these conditions, a separate, post-processing, analysis was conducted with the inclusion of an additional 50% minimum target coverage to assess the quality of the alignments across all applications. Since both target databases contain complete, full-length proteins, this parameter is appropriate. With both, EnTAP reported the highest alignment rate against RefSeq, with Trinotate taking the lead for Swiss-Prot alignments. Overall, Blast2GO annotated fewer sequences than both EnTAP and Trinotate when run against Swiss-Prot and RefSeq. *E. carinata* against Swiss-Prot for Trinotate, Blast2GO, and EnTAP reported annotation rates of: 16.15%, 11.40%, and 15.49%, respectively (Table S7, Figure 3B). Additionally, EnTAP selected a larger percentage of *informative* alignments in two of the three transcriptomes against the Swiss-Prot database. Looking at the results for *P. flexilis*, EnTAP, Trinotate, and Blast2GO selected *informative* alignment percentages for 95.89%, 95.49%, and 94.99%, respectively. In examining the percentage of alignments kept from the original runs, implementing only 50% query coverage, that remained when applying the 50% target coverage, Blast2GO maintained the majority of its alignments. For example, *E. carinata* against Swiss-Prot reported 52.26%, 66.27, and 56.32% for Trinotate, BLAST2GO, and EnTAP, respectively.

Across all three transcriptomes, EnTAP produced slightly higher annotation rates against RefSeq when applying both coverage thresholds (Table S7, Figure 3D). Examining the results from *E. carinata*, EnTAP, Trinotate, and Blast2GO reported 25.11%, 22.74%, and 17.85%, respectively. When including both alignments and gene family/protein domain assignment, EnTAP (55.62%) had a significantly higher annotation rate compared to Trinotate (47.71%) and Blast2GO (5.58%). Examining the percentage of alignments remaining from the original (50% query coverage alignments), Blast2GO had the highest percentage of 71.01%, followed by EnTAP with 65.19%. As before, Trinotate had the lowest percentage of alignments retained after introducing the additional coverage threshold.

#### Trinotate and EnTAP Combined Annotation Methodology

EnTAP is designed to run with at least two databases to provide optimal alignments for non-model organisms. Blast2GO and Trinotate rely on independent database comparisons and do not support mechanisms to merge or select optimal alignments across databases. Trinotate restricts most supplemental information to Swiss-Prot derived data while Blast2GO is able to perform GO mappings across different source databases. Here, we compare a combined run where EnTAP can leverage its alignment algorithm across the same two databases (Swiss-Prot and RefSeq) and Trinotate is executed and combined subsequently. These two applications were compared as they have very similar means of installation and usage compared to Blast2GO, while also having similar homology results. This analysis will include both the 50% query and 50% target coverage for alignments which is the default setting for EnTAP.

#### Trinotate and EnTAP Combined Annotation Results

Three annotation categories were examined between both pipelines: overall annotation rate, Gene Ontology annotation rate, and KEGG annotation rate. EnTAP consistently produced a higher overall annotation compared with Trinotate. Again, overall annotation is described as each sequence receiving either an alignment against the protein databases, or an annotation through HMMER (Trinotate) and EggNOG (EnTAP). *F. hygrometrica* yielded annotation rates of 70.06% and 58.36% for EnTAP and Trinotate, respectively (Figure 5A, Table S12). Gene Ontology and KEGG annotation rates were more varied, with EnTAP yielding higher Gene Ontology assignments for two of the three transcriptomes, while Trinotate consistently produced higher KEGG assignments. The analysis of *P. flexilis* resulted in EnTAP assigning Gene Ontology terms to 39.54% of the sequences, while Trinotate assigned 29.53% of the sequences (Figure 5B). Alternatively, again considering *P. flexilis*, EnTAP and Trinotate assigned KEGG terms to 12.26% and 22.41% of sequences, respectively (Figure 5C). The higher KEGG assignment by Trinotate can be attributed to the pipeline’s reliance on Swiss-Prot.

**Figure 5.**
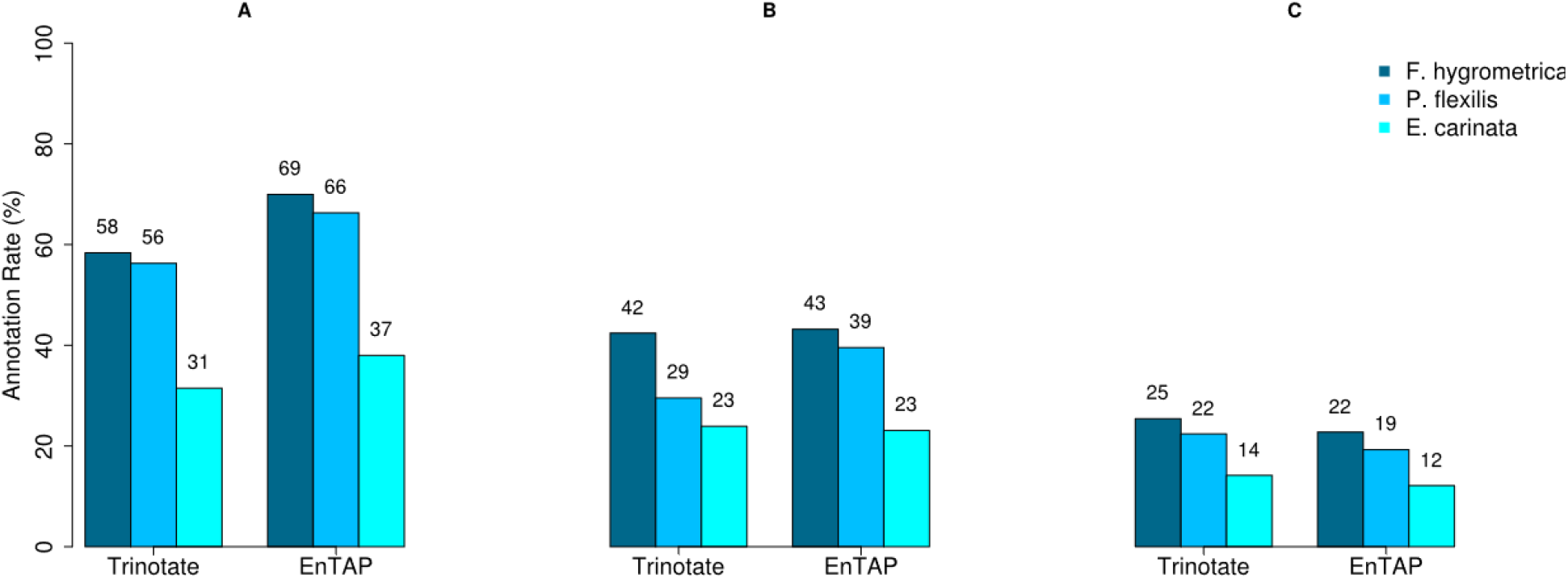
Combined Annotation Results – Trinotate and EnTAP. (A) Overall annotation rate defined as receiving either an alignment against the protein databases, or an annotation through other means. (B) Gene Ontology annotation rate based upon each sequence receiving at least one term assignment. (C) KEGG assignment based upon each sequence receiving at least one assignment.

The selected alignments were analyzed based upon their phylogenetic relevance to the target species, contaminant status, and informativeness. EnTAP utilizes a unique method of removing transcripts when a strong alignment is identified to other organisms associated with contaminants. A similar method is imposed for selecting alignments closer in taxonomic lineage to the species being studied. In all cases, EnTAP filtered out more contaminated alignments than Trinotate across each of the species examined (Table S13, Figure 6B). This trend can be seen when examining the results for *F. hygrometrica*, with Trinotate producing 371 bacteria and 198 fungal contaminants, while EnTAP resulted in 29 bacteria alignments and 33 fungal alignments. EnTAP was able to identify a reliable non-contaminant alignment more times than Trinotate which was naive to the target organism. Furthermore, EnTAP continually selected alignments closer in taxonomic lineage to the target species. A larger distinction can be seen in class alignments, with EnTAP providing more alignments closer to the target species’ class than Trinotate in all transcriptomes analyzed. Considering *F. hygrometrica*, EnTAP selected 5067 alignments in the same Order (*funariales)*, while Trinotate produced 1454 alignments (Figure 6A). Additionally, EnTAP selected fewer informative alignments, based upon a lexicon of terms associated with curated entries when compared to Trinotate (Figure 6C). The lower than expected number of informative alignments with EnTAP is attributed to the introduced bias where we selected alignments with an external script among the two databases for Trinotate. Since Trinotate favors the Swiss-Prot database for assigning additional annotation information, these alignments were selected over their RefSeq counterparts, leading to a higher number of informative alignments and annotation rate compared with EnTAP. Trinotate’s analysis of *F. hygrometrica*, produced an informative percentage of 67.09% compared to 45.26% (EnTAP).

**Figure 6.**
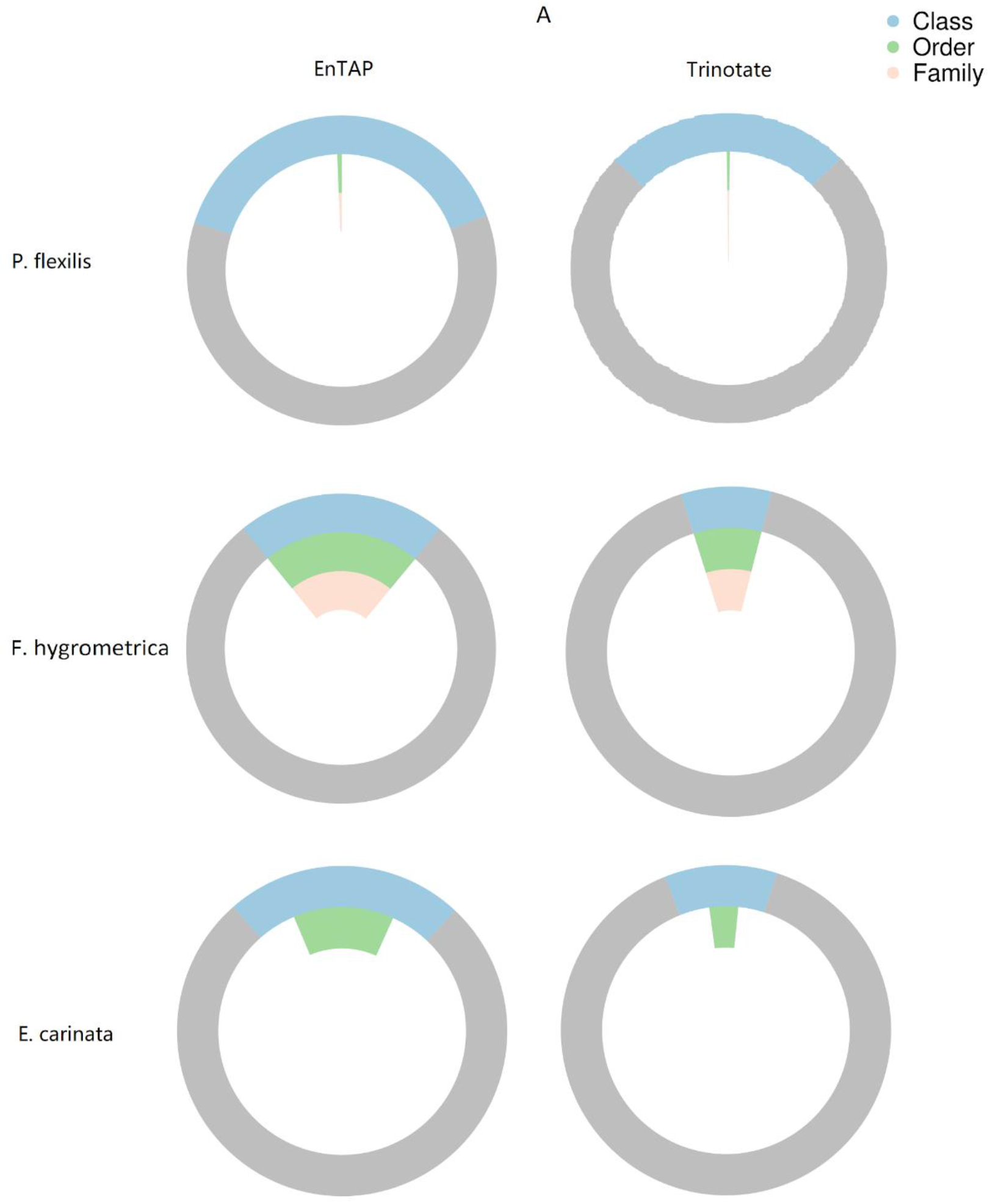
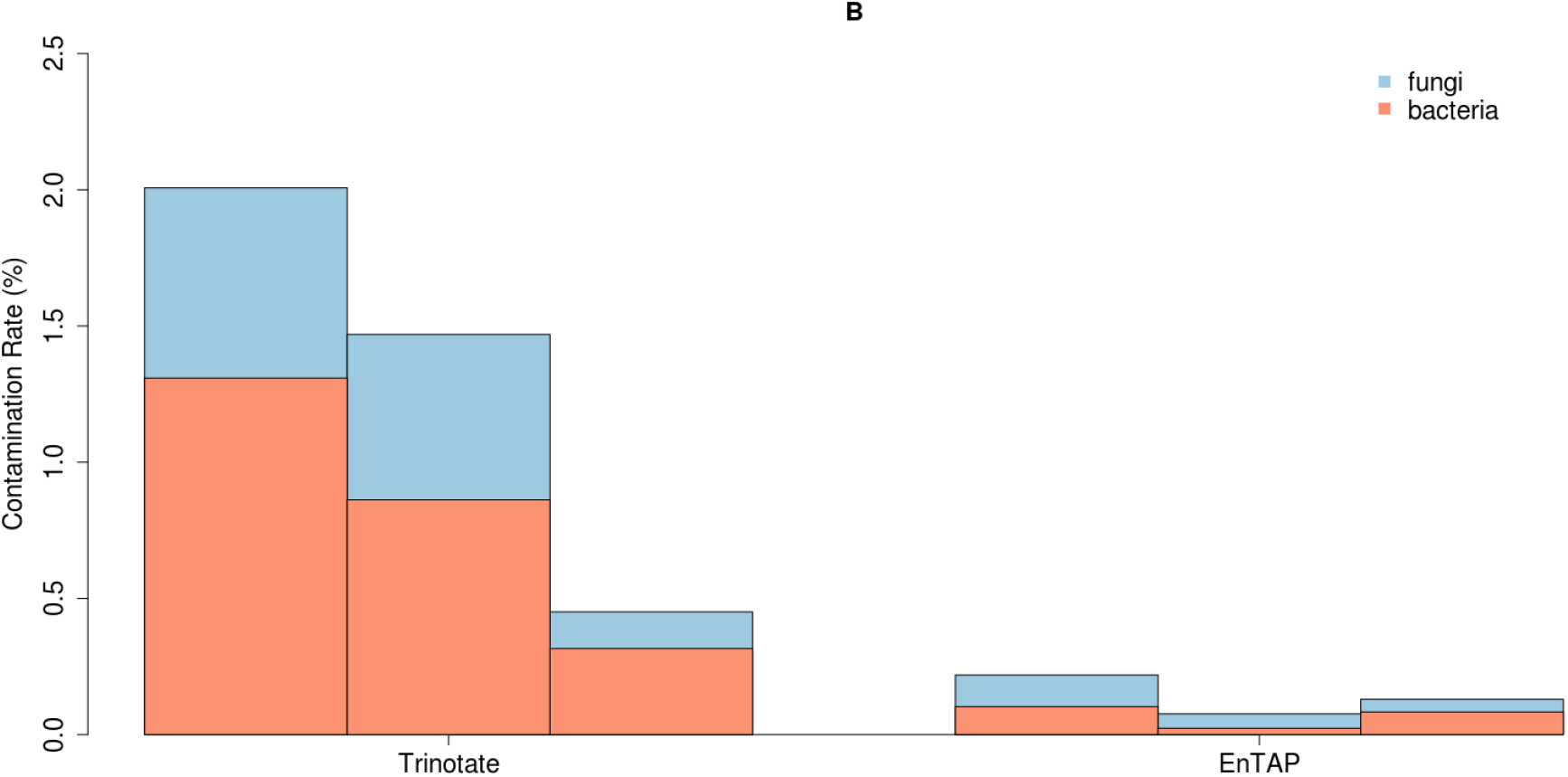
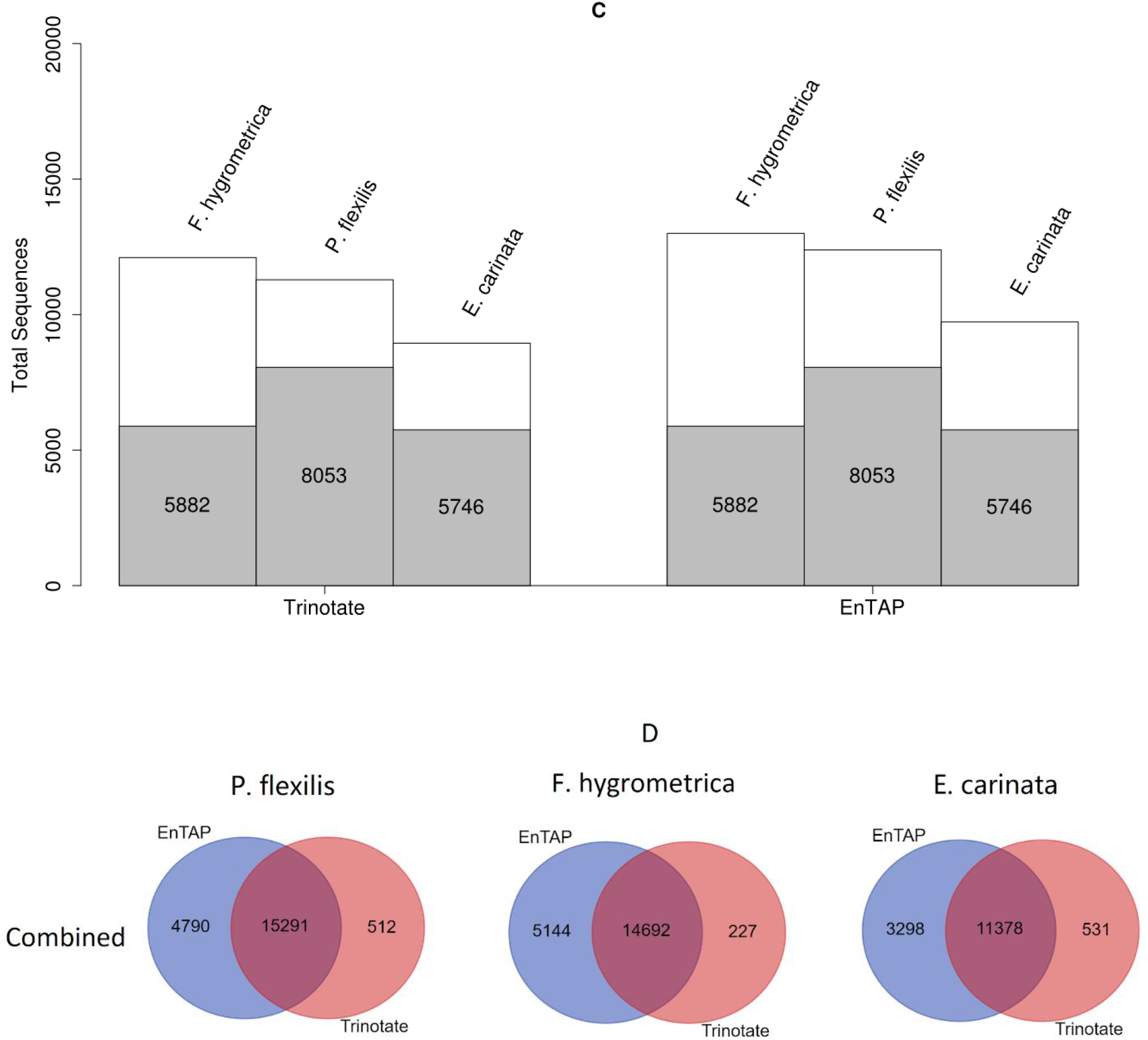
Combined Homology and Annotation Results – EnTAP and Trinotate. Comprehensive analysis of homology, or similarity search, results in phylogenetic relevance to the source species in regards to having the same Class, Order, or Family (A), contaminant detection (B), and informativeness (C). Further analysis to represent sequences annotated between EnTAP and Trinotate in regards to unique annotations or sequences annotated by both pipelines (D).

Additional analyses were conducted to examine the unique sequences annotated among the two pipelines. Results consistently demonstrate that EnTAP annotates more unique sequences when aligning against both Swiss-Prot and RefSeq databases (Figure 6D). When examining the results for *P. flexilis*, EnTAP annotated 4,790 sequences that were not annotated by Trinotate, while Trinotate annotated 512 unique sequences. This disparity can be attributed to the incorporation of EggNOG as opposed to the HMMER means of annotation by Trinotate in addition to the lower quality alignments from Trinotate (BLASTX). The inclusion of 50% query and target coverages eliminated many lower quality alignments from Trinotate, ultimately resulting in a lower annotation rate. Furthermore, frame selection via GeneMarkS-T allowed EnTAP to retain more sequences when compared with Trinotate. In the *P. flexilis* example, both pipelines had an overlapping annotation rate of 15,291.

### Execution Time

In all scenarios, EnTAP’s total execution time (combination of frame selection, similarity searching, protein domain/gene family assignment, and Gene Ontology/KEGG term assignment) was significantly shorter than the other pipelines (Table 2). This analysis took advantage of a high-performance computing cluster for EnTAP and Trinotate, with Blast2GO utilizing a personal computer to emulate the standard user experience. On average, EnTAP completed all steps in under four hours (compared to several days) for all species with moderate sized transcriptomes and modest hardware. This disparity is most apparent with execution against the RefSeq database with an overall EnTAP pipeline execution time varying between 2.6 and 3.55 hours, while Trinotate provided execution times between 409.74 and 748.61 hours, and Blast2GO between 107.76 and 205.05 hours. The large disparities can ultimately be attributed to the faster NCBI BLAST alternative that EnTAP employs, as well as the optimized execution of the entire pathway start to finish. Protein domains or gene family assignments were executed faster in EnTAP compared to Trinotate for *F. hygrometrica* and *P. flexilis*, with approximately 1.8 hours compared to 4 hours, respectively. *E. carinata* gene family/protein domain annotation performed similarly to EnTAP’s with 1.8 hours compared to 2 hours, respectively. Blast2GO’s annotation was the slowest of the three, with *E. carinata*, *P. flexilis*, and *F. hygrometrica* completing at 131.00, 115.41, and 77.05 hours, respectively.

**Table 2.** Pipeline Runtimes.

| Pipeline | UniProt Swiss-Prot |  |  | NCBI RefSeq Complete |  |  | Combined |
| --- | --- | --- | --- | --- | --- | --- | --- |
|  | <u>Trinotate</u> | <u>Blast2GO</u> | <u>EnTAP</u> | <u>Trinotate</u> | <u>Blast2GO</u> | <u>EnTAP</u> | <u>EnTAP</u> |
| ***Funaria hygrometrica (hrs)*** | 16.44 | 81.95 | 1.13 | 409.74 | 107.76 | 2.60 | 2.63 |
| ***Entylia carinata (hrs)*** | 15.22 | 134.23 | 2.07 | 746.68 | 205.05 | 3.90 | 3.73 |
| ***Pinus flexilis (hrs)*** | 17.63 | 119.63 | 1.86 | 748.61 | 148.08 | 3.55 | 3.58 |

### Qualitative Analysis

All three pipelines provide execution through the command line, allowing utilization of high-performance computing clusters to speed up execution time. However, this functionality, along with frame selection, expression analysis, and much more, is blocked behind a paid subscription service for Blast2GO. Both EnTAP and Trinotate provide frame selection services, while only EnTAP provides transcriptome filtering based on expression values. Although it should be noted, Trinotate Web provides some additional functionality for differential expression analysis. Trinotate relies on Swiss-Prot for extracting information but this database is limited to very well curated systems leaving divergent species with minimal information.

All three pipelines attempt to provide an intuitive means of installation and execution with varying levels of success for the average user. The installation process for the Java-based version of Blast2GO is, naturally, the simplest as it is a standalone desktop application with very few steps to fully install the software. EnTAP and Trinotate both have fairly similar installation processes based on command line exeuction. However, users may run into some difficulties when installing the multiple Trinotate dependencies and setting up the SQL database.

The usability and flexibility of the pipelines varies greatly in regards to features and subscriptions status. Database variability is limited in Trinotate and Blast2GO, while EnTAP permits any database with the minor exception that the sequence header must be formatted in NCBI or EBI formats for full taxonomic or contaminate filtering. As discussed, Trinotate has a heavy reliance on the Swiss-Prot database where the majority of the Gene Ontology and all of the pathway information is derived from. Blast2GO allows a variety of databases to be created as long as the subscription service is purchased. However, neither platform can integrate across different database sources. EnTAP is the only pipeline that incorporates an optimal alignment algorithm to select and filter alignments across a variety of databases. As a result, executing EnTAP against several databases will produce a single optimal alignment for each sequence filtered by contaminant and taxonomic status. This is beneficial for non-model systems since source databases vary by levels of curation and size.

## DISCUSSION

Non-model eukaryotic annotation presents several challenges associated with processing time, integration of existing genetic resources, and annotation quality that are not yet fully resolved with existing pipelines. We present EnTAP, a novel method of open-source transcriptome annotation improves upon the existing solutions.

We performed a comprehensive analysis between EnTAP and two of the most widely used pipelines, Trinotate and Blast2GO Pro. Overall annotation and homology rates, methods of transcriptome filtering, and available pipeline features were discussed. From the three independent eukaryotic transcriptomes analyzed, both pooled and non-pooled, EnTAP’s method of frame selection produced more complete genes compared with Trinotate, leading to a higher quality annotation downstream. Homology results against NCBI’s RefSeq database, present highest rates of alignment for EnTAP when considering a higher quality of alignment and similar rates between Trinotate against Swiss-Prot. Blast2GO had the lowest rate of overall alignment, but the highest percentage of quality alignments. The overall annotation rate, quantified as an alignment through homology or through HMMER or EggNOG without quality thresholds, varied between the pipelines with EnTAP having the highest rates against Swiss-Prot, yet slightly lower rates compared with Trinotate against RefSeq. With all transcriptomes, EnTAP had the fastest execution, up to 50 times faster than Blast2GO and even faster than Trinotate. A combined analysis with both databases and the quality thresholds of 50% query and 50% target coverage provided the highest overall annotation and alignment rate with EnTAP.

EnTAP is a versatile, fast, and accurate non-model annotation pipeline that provides a complete annotation approach, beginning with optional transcriptome filtering through frame selection or expression analysis, and concluding with annotation through homology and gene family assignment. By leveraging accompanying software packages and its unique methods of optimal alignment selection and contaminant filtering, EnTAP can provide a personalized, comprehensive, and efficient annotation making it a viable alternative to existing solutions.

## MATERIALS AND METHODS

### Data and Database Acquisition

A total of three non-model, Illumina HiSeq paired-end sequenced and *de novo* assembled transcriptomes were acquired for evaluating EnTAP. All organisms were assembled *de novo* with Trinity v2.06 (*F. hygrometrica* and *P. flexilis)* and v2.2 (*E. carinata)*. The three species chosen were the *Entylia carinata* (keeled treehopper), *Funaria hygrometrica* (cord-moss), and *Pinus flexilis* (limber pine) providing a varied taxonomic range for comparison.

The transcriptomes ranged in size from 28,350 to 38,640 transcripts. The libraries represent single genotype RNA extractions in the case of *P. flexilis* and *F. hygrometrica*, and pooled libraries for *E. carinata*. The raw reads (SRA) and assembled data (TSA) are available via NCBI (PRJNA415461, PRJNA421369, PRJNA254339).

The input sequence sets were evaluated against the same public databases. Swiss-Prot (accessed September 26th, 2017) and NCBI RefSeq Complete (v84). Additionally, the NCBI Taxonomic Database (accessed July 21st, 2017) [23] was used for evaluation of contaminants and taxonomic relevance of alignments. The Gene Ontology database was accessed for additional description and categorical information (accessed August 20th, 2017). The EggNOG DIAMOND configured database was accessed through EggNOG-mapper (accessed August 20th, 2017). Blast2GO databases were accessed through the CloudBlast service for pro users (accessed October 2017).

### EnTAP implementation

EnTAP was developed in C/C++ language and is designed for a Unix-based environment. The Boost C++ Libraries (1.50 or later) [26] provide a reliable means of generic typing. Cereal [27] provides C++ serialization methods for rapid accession of mapping information, while the TCLAP library [28] was utilized as a simple command line parser. CMake (2.8 or later) [29] made for an intuitive means of dependency verification and Makefile generation. A C++ interface for POSIX process control and error and output piping was provided through the library PStreams (0.8.1 or later) [30]. Additional multi-threaded file parsing was accomplished through the use of “Fast C++ CSV Parser.” Python (2.7.12 or later module) [31] allowed for graphical representations of the data (through “matplotlib” [32] module), SQLITE lookups of the EggNOG database (through “sqlite3”[33]), and querying the NCBI Taxonomic Database. An additional SQLITE [34] interface is included in the EnTAP repository for accession of the EggNOG databases outside of Python. A compiler that supports C++11 features is required for compilation. EnTAP (beta version 0.7.4) supports the following accompanying pipeline software: RSEM (versions 1.3.0), GeneMarkS-T (version 5.1), DIAMOND (version 0.8.31), and EggNOG-Emapper (version 0.7.4.1-beta).

EnTAP execution is divided into two stages: *configuration* and downloading of pertinent databases, and *execution* of the main annotation pipeline. All exceptions that interfere with pipeline execution, in either stage, are handled as fatal errors with distinct error messages provided from either the pipeline software or EnTAP to easily identify the source of the failure. Additionally, a debug file is updated at every stage.

*Configuration* of EnTAP must be run once to download and index the NCBI Taxonomic Database [25], Gene Ontology Database [35], EggNOG databases [36], and an optional number of DIAMOND databases to remove the requirement of an Internet connection during the annotation stage. These databases are not included in the EnTAP repository and must be downloaded separately. The NCBI Taxonomic Database and Gene Ontology Database are pre-formatted and hosted for EnTAP to download. Python is utilized to query the NCBI Taxonomic Database for all entries. Entries containing lineage and NCBI Accession IDs are formatted into a serialized hash map that can be read back into memory upon *execution*. A comparable method is used to retrieve, extract, and serialize a map of Gene Ontology terms with pertinent term and accession information derived from the Gene Ontology Consortium [19]. This method is incorporated to sidestep an oftentimes cumbersome full SQL database installation while maintaining a great deal of information. Additionally, the EggNOG databases are downloaded through the EggNOG-mapper software while DIAMOND is incorporated to index FASTA protein databases into a compatible format for DIAMOND. Users have the option to specify output directories and execution paths of the software involved. Upon completion of *configuration*, *execution* can be performed.

*Execution* requires, from the user, a single multi-FASTA transcriptome file and specification of up to five DIAMOND formatted databases. EnTAP is ignorant of the assembler used and will provide an annotation regardless of annotation software. It will optionally accept an un-gapped alignment file in SAM or BAM format for use in *transcriptome filtering* using a default FPKM (fragments per kilobase million) cutoff of 0.5 that can be modified by the user. *Transcriptome filtering* is implemented with RSEM [13] and followed by GeneMarkS-T [37] spawned as child processes through the PStreams library. EnTAP utilizes Python with the Matplotlib module to generate graphical analyses of the data by passing information derived through RSEM and GeneMarkS-T results. Standard error and standard output are directed to EnTAP and written for the user to their respective files.

The *transcriptome annotation* stage of *execution* can be customized by the user. Homology, or similarity searching against reference databases, allows for specifying target and query coverage cutoffs (default value: 50%), E-value (default value: 1E-5), contaminant and target taxon information for phylogenetic filtering, and an additional list of “uninformative” terms (default listing: *conserved, predicted, unnamed, hypothetical, putative, unidentified, uncharacterized, unknown, uncultured, uninformative*). EnTAP utilizes E-value, contaminant status, coverage, taxonomic relevance, and informativeness of the description to select the most optimal hit (Figure S1). A taxonomic score is developed based upon the taxonomic relevance of the hit to the target species and the informativeness of the alignment.

Most notable differences in EnTAP’s selection of the optimal alignments compared to other pipelines is seen in alignments that are very similar in quality, or E-value and coverage. The selection process begins with a comparison of both parameters derived from the information generated from DIAMOND. If the values of each alignment are within a predetermined range, they will continue to the next decision. However, when comparing against the same database, if one alignment is superior in terms of E-value or coverage, it will be selected. Due to varying E-values across databases, coverage is utilized when comparing different databases. This is incorporated to ensure the higher quality hits are not removed due to the decision processes that follow. Assuming alignments are within a similar quality range, EnTAP will evaluate contaminant status. This is done by mapping the species information derived from either an EMBL or NCBI formatted fasta header to the previously downloaded NCBI Taxonomic Database. Through this quick accession, the lineage is determined and compared with the taxonomic contaminants provided by the user. From here, the contaminant is removed and the non-contaminant remains. If both alignments are considered contaminants or non-contaminants, a combined analysis of taxonomic relevance and informativeness will follow. Using the previously mapped phylogenetic lineage information, EnTAP compares this to the user provided “target species,” or the transcriptome origin species, and determines a score based upon its taxonomic similarity to the alignment. The factor of informativeness is then compounded upon this result, with a more informative alignment weighted above an uninformative one. An informative rating, or level of curation, will be the deciding factor between redundant annotations across separate databases.

Gene family assignment is made available by the EggNOG databases and EggNOG-mapper as a means of accessing them. Through EggNOG-mapper, DIAMOND is again leveraged to align transcripts to the EggNOG database. Optimal alignments are selected purely based upon E-value and lookups of the EggNOG SQL database are performed through the “sqlite3” module of Python. Further Gene Ontology, pathway, and functional information is derived from the SQL database through lookups performed by EnTAP. Gene Ontology terms are mapped to the Gene Ontology database previously indexed to determine additional description and categorization. Python’s Matplotlib module is again utilized to produce histogram graphical representations of Gene Ontology categorical (molecular function, cellular component, and biological process) and level distributions, gene family taxonomic distributions, and pathway information piped from EnTAP to the Python plotting script.

### Evaluation

EnTAP (beta v0.7.4) was compared against Trinotate (v3.0.2) and Blast2GO Pro (v4.1.9). Trinotate and EnTAP were installed on a compute cluster and executed with 8 threads on AMD Opteron CPU with 128 GB RAM. Blast2GO Pro was installed on a personal computer running Windows 10 with an Intel Core i7-6700k running at 4.20 GHz and 16 GB RAM.

#### Transcriptome Filtering

Pre-processing of frame was enabled via GeneMarkS-T (EnTAP, GeneMarkS-T v5.1) and Transdecoder (Trinotate, Transcoder v3.0.0). Default parameters were used for frame selection in both cases. A limited expression filtering analysis was performed for EnTAP following frame selection against *E. carinata* and *F. hygrometrica* transcriptomes with RSEM (default parameters with an FPKM cutoff of 0.5 post-processing, v1.3.0). The runs without RSEM were used for direct comparison against Trinotate and Blast2GO.

#### Transcriptome Annotation

Sequences filtered through frame selection or expression analysis were removed from annotation for EnTAP and passed to the sequence similarity search stage. Trinotate maintains the sequences in which a frame was not found, performing BLASTX. DIAMOND (EnTAP, DIAMOND v0.8.3.1) and NCBI BLAST+ (Trinotate, BLAST+ v2.4.0), and CloudBlast (Blast2GO) were implemented with the same parameters (E-value: 1E-5, query coverage: 50%, total alignments: 3). All applications were executed with the same two public databases (UniProt Swiss-Prot and NCBI RefSeq). Optimal alignments were calculated with EnTAP’s custom method (Figure S1) and the best hit was selected via E-value score for Trinotate with alignments from BLASTP favored over BLASTX. Blast2GO selections of the best hit were primarily based upon the E-value. Contaminant filtering was employed in EnTAP for the following groups: *bacterial* and *fungal*. These designations could not be parameterized for Blast2GO or Trinotate so post-filtering approaches were used to evaluate the number of assignments back to these categories. The NCBI Taxonomy database is used as the reference for this definition by determining the lineage from the origin species contained within the headers of the supported databases. Additionally, this mapping was used to determine the taxonomic relevance of alignments. Taxonomic relevance calculations were based upon the alignment species’ relevance to the target species in relation to genus, family, order, and class. Analysis of *E. carinata* used *membracidae*, *hemiptera*, and *insecta* as family, order, and class categories. Analysis of *P. flexilis* used *pinaceae*, *pinales*, and *spermatophyta* as family, order, and class categories. Analysis of *F. hygrometrica* used *funariaceae*, *funariales*, and *bryopsida* as family, order, and class categories. Further annotation was performed through EggNOG-mapper (EnTAP, v0.8.0-beta), InterProScan (Blast2GO), and HMMER (Trinotate, v3.1b2) to assign Gene Ontology terms and protein domains.

## ACKNOWLEDGEMENTS

We thank the members of the Plant Computational Genomic Lab and Computational Biology Core at the University of Connecticut including Dr. Uzay Sezen, Alex Trouern-Trend, Sumaira Zaman, Dr. Neranjan Perera, Jeff Lary, and Dr. Vijender Singh for their continued support and insight during the development process. We also thank Dr. Claudio Casola of the University of Texas A&M University, Erik Visser of the University of Pretoria, Susan McEvoy of the Oregon State University, and Dr. Kevin Brown and Dr. Rachel O’Neill of the University of Connecticut who have provided a variety of datasets and suggestions to aid in development of the pipeline.

## DATA ACCESSIBILITY

EnTAP is available from https://gitlab.com/EnTAP/EnTAP under the open source license GNU General Public License v3.0. Detailed documentation is available here: http://entap.readthedocs.io/en/latest/. The version of the source code used in the manuscript is https://gitlab.com/EnTAP/EnTAP/tags/v0.7.4-beta. All datasets analyzed in this study, are available from NCBI as PRJNA415461, PRJNA421369, and PRJNA254339.

## AUTHORS’ CONTRIBUTIONS

AJH, SG, RP, and JLW conceived and conducted the study; AJH developed the EnTAP algorithm; AJH, MX, and JLW designed follow-up experiments, including EnTAP testing; JLW supervised research; AJH, MX, and JLW wrote the manuscript, with input from other authors. NR provided the data and metadata for *Funaria hygrometrica*. CRF provided the data and metadata for *Entylia carinata*. JBM provided the data and metadata for *Pinus flexilis*.

All authors read and approved the final manuscript.

