## Supplemental Data for "EnTAP: Bringing Faster and Smarter Functional Annotation to Non-Model Eukaryotic Transcriptomes"

**Table S1**. *Funaria hygrometrica* Frame Selection Results

| | Trinotate (Transdecoder) | EnTAP (GeneMarkS-T) |
|---|---|---|
| **Average Sequence Length (bp)** | 808.79 | 761.39 |
| **N50** | 975 | 927 |
| **No Frame Detected** | 9017 | 6635 |
| **Frame Detected** | 19333 | 21715 |
| **Partial 5'** | 5347 | 4839 |
| **Partial 3'** | 2122 | 2719 |
| **Complete** | 7213 | 9695 |
| **Internal** | 5428 | 4462 |

**Table S2.** *Entylia carinata* Frame Selection Results

| | Trinotate (Transdecoder) | EnTAP (GeneMarkS-T) |
|---|---|---|
| **Average Sequence Length (bp)** | 1012.49 | 946.33 |
| **N50** | 1302 | 1245 |
| **No Frame Detected** | 23549 | 20256 |
| **Frame Detected** | 15091 | 18384 |
| **Partial 5'** | 4179 | 3748 |
| **Partial 3'** | 1576 | 2355 |
| **Complete** | 5724 | 7467 |
| **Internal** | 5539 | 4814 |

**Table S3.** *Pinus flexilis* Frame Selection Results

| | Trinotate (Transdecoder) | EnTAP (GeneMarkS-T) |
|---|---|---|
| **Average Sequence Length (bp)** | 835.59 | 815.40 |
| **N50** | 1011 | 984 |
| **No Frame Detected** | 8586 | 6595 |
| **Frame Detected** | 21705 | 23696 |

| | | |
|---|---|---|
| **Partial 5’** | 4709 | 5543 |
| **Partial 3’** | 3580 | 3847 |
| **Complete** | 5900 | 7580 |
| **Internal** | 7516 | 6726 |

**Table S4.** *Funaria hygrometrica* Homology Results

| | **UniProt Swiss-Prot** | | | **NCBI RefSeq Complete** | | |
|---|---|---|---|---|---|---|
| **Pipeline** | Trinotate | Blast2GO | EnTAP | Trinotate | Blast2GO | EnTAP |
| **Alignments** | 14726 | 8995 | 13540 | 20763 | 13888 | 19468 |
| **Percentage of Transcriptome (%)** | 51.94 | 31.73 | 47.76 | 73.24 | 48.99 | 68.67 |
| **Fungi Alignments (%)** | 2.44 | 2.22 | 1.63 | 0.37 | 0.36 | 0.28 |
| **Bacteria Alignments (%)** | 3.66 | 5.67 | 3.38 | 0.18 | 0.40 | 0.02 |
| **Informative Percentage (%)** | 96.00 | 95.49 | 96.43 | 6.79 | 5.73 | 8.19 |
| **Phylogeny Genus Alignments (%)** | 0.00 | 0.00 | 0.00 | 0.00 | 0.00 | 0.00 |
| **Phylogeny Family Alignments (%)** | 0.76 | 1.00 | 0.82 | 92.65 | 90.31 | 92.09 |
| **Phylogeny Order Alignments (%)** | 0.76 | 1.00 | 0.82 | 92.65 | 90.31 | 92.09 |
| **Phylogeny Class Alignments (%)** | 0.88 | 1.01 | 0.92 | 92.66 | 90.32 | 92.10 |

**Table S5.** *Entylia carinata* Homology Results

| | **UniProt Swiss-Prot** | | | **NCBI RefSeq Complete** | | |
|---|---|---|---|---|---|---|
| **Pipeline** | Trinotate | Blast2GO | EnTAP | Trinotate | Blast2GO | EnTAP |
| **Alignments** | 11938 | 6649 | 10628 | 16442 | 9715 | 14796 |
| **Percentage of Transcriptome (%)** | 30.89 | 17.21 | 27.51 | 42.56 | 25.14 | 38.51 |
| **Fungi Alignments (%)** | 0.69 | 0.99 | 0.48 | 0.29 | 0.28 | 0.23 |
| **Bacteria Alignments (%)** | 0.93 | 1.03 | 0.72 | 1.51 | 1.43 | 0.28 |
| **Informative Percentage (%)** | 97.70 | 99.00 | 97.76 | 42.49 | 5.85 | 57.87 |
| **Phylogeny Genus Alignments (%)** | 0.00 | 0.00 | 0.00 | 0.04 | 0.07 | 0.00 |
| **Phylogeny Family Alignments (%)** | 0.00 | 0.00 | 0.00 | 0.04 | 0.07 | 0.00 |
| **Phylogeny Order Alignments (%)** | 0.28 | 0.45 | 0.38 | 54.68 | 47.44 | 60.04 |
| **Phylogeny Class Alignments (%)** | 32.14 | 31.19 | 34.92 | 92.17 | 89.06 | 95.69 |

**Table S6.** *Pinus flexilis* Homology Results

| | UniProt Swiss-Prot | | | NCBI RefSeq Complete | | |
|---|---|---|---|---|---|---|
| **Pipeline** | Trinotate | Blast2GO | EnTAP | Trinotate | Blast2GO | EnTAP |
| **Alignments** | 16718 | 9209 | 15357 | 21182 | 13105 | 19823 |
| **Percentage of Transcriptome (%)** | 55.19 | 30.40 | 50.69 | 69.93 | 43.26 | 65.44 |
| **Fungi Alignments (%)** | 1.81 | 2.66 | 1.31 | 0.57 | 0.83 | 0.21 |
| **Bacteria Alignments (%)** | 2.12 | 3.67 | 1.92 | 1.32 | 2.49 | 0.01 |
| **Informative Percentage (%)** | 95.69 | 94.47 | 96.31 | 33.78 | 27.29 | 63.04 |
| **Phylogeny Genus Alignments (%)** | 1.03 | 1.25 | 1.31 | 0.16 | 0.18 | 0.17 |
| **Phylogeny Family Alignments (%)** | 1.68 | 2.28 | 1.93 | 0.16 | 0.21 | 0.17 |
| **Phylogeny Order Alignments (%)** | 1.68 | 2.28 | 1.93 | 0.16 | 0.21 | 0.17 |
| **Phylogeny Class Alignments (%)** | 86.42 | 83.42 | 87.41 | 90.11 | 86.84 | 93.85 |

**Table S7.** Homology Results (50% Query and 50% Target Coverage)

| | UniProt Swiss-Prot | | | NCBI RefSeq Complete | | |
|---|---|---|---|---|---|---|
| **Pipeline** | Trinotate | Blast2GO | EnTAP | Trinotate | Blast2GO | EnTAP |
| | | | | | | ***Funaria hygrometrica*** |
| **Alignments** | 8359 | 5973 | 8090 | 11883 | 9561 | 12965 |
| **Percentage of Transcriptome (%)** | 29.49 | 21.07 | 28.54 | 41.92 | 33.72 | 45.73 |
| **Percentage of Informative Alignments (%)** | 95.81 | 95.78 | 96.18 | 6.03 | 6.29 | 12.31 |
| **Percentage of Quality Alignments (%)** | 56.76 | 66.40 | 59.75 | 57.23 | 68.84 | 66.59 |
| | | | | | | ***Entylia carinata*** |
| **Alignments** | 6239 | 4403 | 5986 | 8786 | 6899 | 9703 |
| **Percentage of Transcriptome (%)** | 16.15 | 11.40 | 15.49 | 22.74 | 17.85 | 25.11 |
| **Percentage of Informative Alignments (%)** | 98.06 | 97.75 | 98.03 | 47.71 | 5.58 | 55.62 |
| **Percentage of Quality Alignments (%)** | 52.26 | 66.27 | 56.32 | 53.44 | 71.01 | 65.19 |
| | | | | | | ***Pinus flexilis*** |
| **Alignments** | 7980 | 6673 | 7800 | 11065 | 10014 | 12350 |
| **Percentage of Transcriptome (%)** | 26.34 | 22.03 | 25.75 | 36.53 | 33.06 | 40.77 |
| **Percentage of Informative** | 95.49 | 94.99 | 95.89 | 33.77 | 28.72 | 58.66 |

| | | | | | | |
|---|---|---|---|---|---|---|
| **Alignments (%)** | | | | | | |
| **Percentage of Quality Alignments (%)** | 47.73 | 72.46 | 50.79 | 52.24 | 76.41 | 62.30 |

**Table S8.** *Funaria hygrometrica* Annotation Results

| | **UniProt Swiss-Prot** | | | **NCBI RefSeq Complete** | | |
|---|---|---|---|---|---|---|
| **Pipeline** | Trinotate | Blast2GO | EnTAP | Trinotate | Blast2GO | EnTAP |
| **Overall Annotation Percentage (%)** | 51.94 | 53.11 | 68.14 | 73.23 | 61.81 | 71.74 |
| **Overall Annotation without Contaminants (%)** | 48.78 | 50.86 | 65.75 | 72.83 | 61.44 | 71.53 |
| **Sequences Annotated With At Least One Gene Ontology Term (%)** | 50.74 | 42.99 | 43.19 | 31.71 | 46.19 | 43.19 |
| **Sequences Annotated With At Least One Pathway Term (%)** | 45.51 | 9.45 | 22.75 | 0.00 | 9.92 | 22.75 |

**Table S9.** *Entylia carinata* Annotation Results

| | **UniProt Swiss-Prot** | | | **NCBI RefSeq Complete** | | |
|---|---|---|---|---|---|---|
| **Pipeline** | Trinotate | Blast2GO | EnTAP | Trinotate | Blast2GO | EnTAP |
| **Overall Annotation Percentage (%)** | 30.89 | 31.85 | 37.53 | 42.55 | 35.31 | 39.47 |
| **Overall Annotation without Contaminants (%)** | 30.39 | 31.49 | 37.19 | 41.78 | 34.88 | 39.27 |
| **Sequences Annotated With At Least One Gene Ontology Term (%)** | 30.08 | 24.45 | 23.08 | 17.98 | 20.14 | 23.08 |
| **Sequences Annotated With At Least One Pathway Term (%)** | 26.49 | 4.50 | 12.13 | 0.00 | 0.88 | 12.13 |

**Table S10.** *Pinus flexilis* Annotation Results

| | **UniProt Swiss-Prot** | | | **NCBI RefSeq Complete** | | |
|---|---|---|---|---|---|---|
| **Pipeline** | Trinotate | Blast2GO | EnTAP | Trinotate | Blast2GO | EnTAP |
| **Overall Annotation Percentage (%)** | 55.19 | 55.53 | 68.18 | 69.93 | 60.01 | 67.23 |
| **Overall Annotation without Contaminants (%)** | 53.03 | 53.61 | 64.54 | 68.61 | 59.55 | 67.09 |
| **Sequences Annotated With At Least One Gene Ontology Term (%)** | 54.12 | 43.54 | 39.54 | 31.52 | 41.34 | 39.54 |

| | | | | | | |
|---|---|---|---|---|---|---|
| **Sequences Annotated With At Least One Pathway Term (%)** | 48.27 | 9.59 | 19.26 | 0.00 | 5.05 | 19.26 |

**Table S11.** Detailed Execution Times

| | **UniProt Swiss-Prot** | | | **NCBI RefSeq Complete** | | |
|---|---|---|---|---|---|---|
| **Pipeline** | Trinotate | Blast2GO | EnTAP | Trinotate | Blast2GO | EnTAP |
| ***Funaria hygrometrica*** | | | | | | |
| **Frame Selection (hrs)** | 0.51 | N/A | 0.02 | 0.51 | N/A | 0.02 |
| **Similarity Search / Homology (hrs)** | 12.05 | 4.90 | 0.03 | 405.35 | 30.76 | 1.5 |
| **Protein Domain Annotation (hrs)** | 3.88 | 77.05 | 1.08 | 3.88 | 77.00 | 1.08 |
| **Overall Execution Time (hrs)** | 16.44 | 81.95 | 1.13 | 409.74 | 107.76 | 2.6 |
| ***Entylia carinata*** | | | | | | |
| **Frame Selection (hrs)** | 0.88 | N/A | 0.04 | 0.88 | N/A | 0.04 |
| **Similarity Search / Homology (hrs)** | 12.54 | 3.23 | 0.03 | 744.00 | 74.05 | 1.66 |
| **Protein Domain Annotation (hrs)** | 1.80 | 131.00 | 2.00 | 1.80 | 131.00 | 2.00 |
| **Overall Execution Time (hrs)** | 15.22 | 134.23 | 2.07 | 746.68 | 205.05 | 3.90 |
| ***Pinus flexilis*** | | | | | | |
| **Frame Selection (hrs)** | 0.81 | N/A | 0.03 | 0.81 | N/A | 0.03 |
| **Similarity Search / Homology (hrs)** | 12.72 | 4.22 | 0.025 | 744 | 32.67 | 1.72 |
| **Protein Domain Annotation (hrs)** | 4.1 | 115.41 | 1.8 | 3.8 | 115.41 | 1.8 |
| **Overall Execution Time (hrs)** | 17.63 | 119.63 | 1.86 | 748.61 | 148.08 | 3.55 |

**Table S12.** EnTAP and Trinotate Combined Annotation Results

| | **Combined (UniProt Swiss-Prot and NCBI RefSeq Complete)** | |
|---|---|---|
| **Pipeline** | Trinotate | EnTAP |
| ***Funaria hygrometrica*** | | |
| **Overall Annotation Percentage (%)** | 58.36 | 69.97 |
| **Sequences Annotated With At Least One Gene Ontology Term (%)** | 42.47 | 43.19 |
| **Sequences Annotated With At** | 25.41 | 22.75 |

| | | |
|---|---|---|
| **Least One Pathway Term (%)** | | |
| | | ***Entylia carinata*** |
| **Overall Annotation Percentage (%)** | 31.47 | 37.98 |
| **Sequences Annotated With At Least One Gene Ontology Term (%)** | 23.91 | 23.08 |
| **Sequences Annotated With At Least One Pathway Term (%)** | 14.16 | 12.13 |
| | | ***Pinus flexilis*** |
| **Overall Annotation Percentage (%)** | 56.28 | 66.29 |
| **Sequences Annotated With At Least One Gene Ontology Term (%)** | 29.53 | 39.54 |
| **Sequences Annotated With At Least One Pathway Term (%)** | 22.41 | 19.26 |

**Table S13.** EnTAP and Trinotate Combined Homology Results

| | **Combined (UniProt Swiss-Prot and NCBI RefSeq Complete)** | |
|---|---|---|
| **Pipeline** | Trinotate | EnTAP |
| | | ***Funaria hygrometrica*** |
| **Alignments** | 12105 | 12997 |
| **Fungi Alignments (%)** | 1.64 | 0.25 |
| **Bacteria Alignments (%)** | 3.06 | 0.22 |
| **Informative Percentage (%)** | 67.09 | 45.26 |
| **Phylogeny Genus Alignments (%)** | 0.00 | 0.00 |
| **Phylogeny Family Alignments (%)** | 28.33 | 47.70 |
| **Phylogeny Order Alignments (%)** | 28.33 | 47.70 |
| **Phylogeny Class Alignments (%)** | 28.34 | 47.72 |
| | | ***Entylia carinata*** |
| **Alignments** | 8944 | 9731 |
| **Fungi Alignments (%)** | 0.58 | 0.18 |
| **Bacteria Alignments (%)** | 1.36 | 0.33 |
| **Informative Percentage (%)** | 76.74 | 59.05 |
| **Phylogeny Genus Alignments (%)** | 0.00 | 0.00 |

| | | |
|---|---|---|
| **Phylogeny Family Alignments (%)** | 0.00 | 0.00 |
| **Phylogeny Order Alignments (%)** | 16.26 | 52.07 |
| **Phylogeny Class Alignments (%)** | 47.67 | 93.17 |
| | | ***Pinus flexilis*** |
| **Alignments** | 11285 | 12386 |
| **Fungi Alignments (%)** | 1.63 | 0.13 |
| **Bacteria Alignments (%)** | 2.31 | 0.01 |
| **Informative Percentage (%)** | 73.02 | 65.02 |
| **Phylogeny Genus Alignments (%)** | 0.92 | 1.03 |
| **Phylogeny Family Alignments (%)** | 1.59 | 1.54 |
| **Phylogeny Order Alignments (%)** | 1.59 | 1.54 |
| **Phylogeny Class Alignments (%)** | 86.43 | 96.02 |

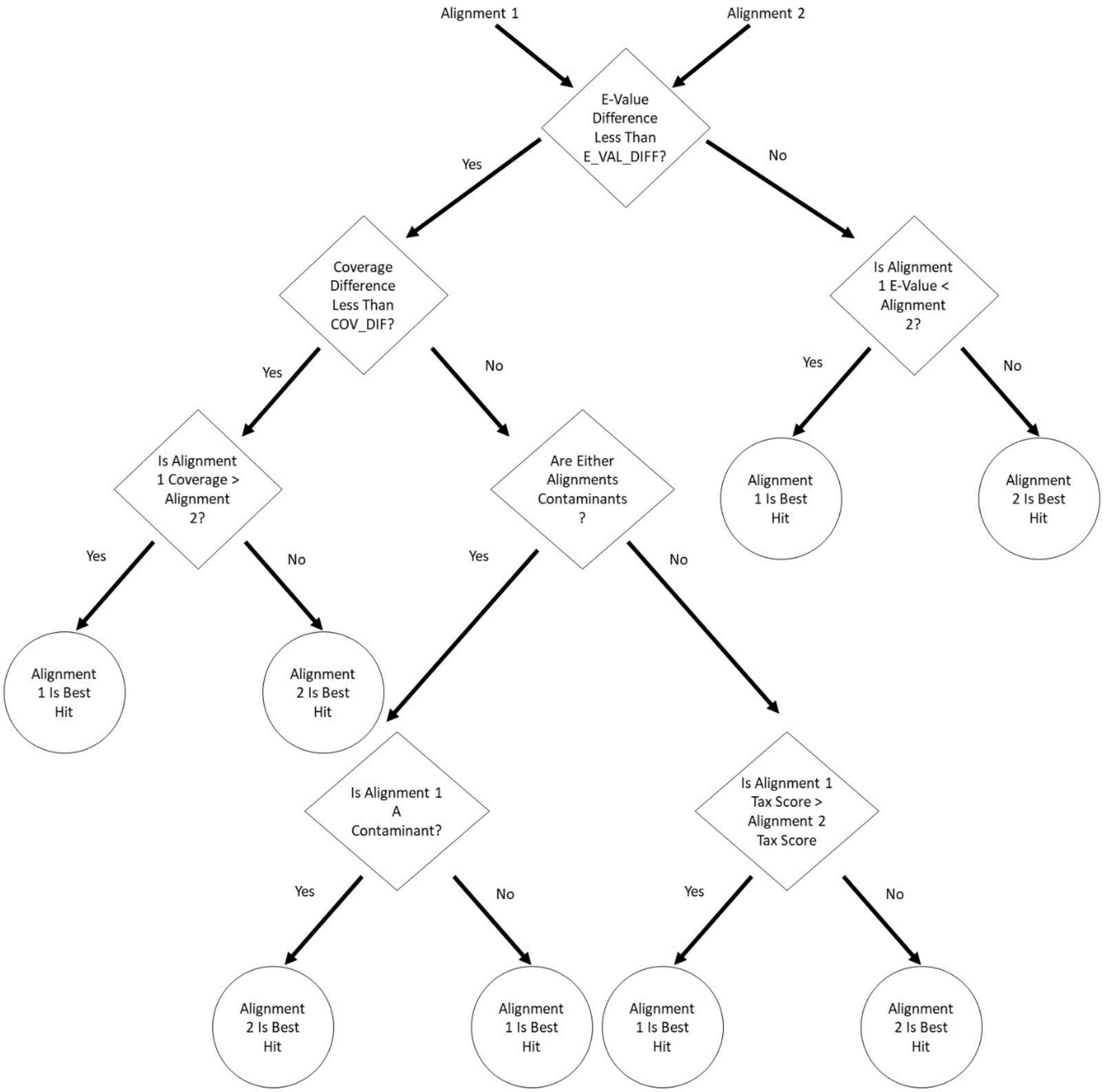


**Figure S1.** EnTAP Optimal Alignment Selection Process
